# Paw switching with lateralized cholinergic modulation

**DOI:** 10.1101/2025.10.17.683000

**Authors:** Kazuki Okamoto, Yasuhiro R. Tanaka, Shigeki Kato, Shu Xie, Guochuan Li, Kazuto Kobayashi, Masato Koike, Yulong Li, Hiroyuki Hioki

**Author notes:** **Correspondence**: Kazuki Okamoto, Ph.D.,; Hiroyuki Hioki, M.D., Ph.D. **Current affiliation:** Department of Cellular Neuropathology, Brain Research Institute, Niigata University, Niigata 951-8585, Japan (Kazuki Okamoto).

## Abstract

Motor preferences, such as handedness, reflect fundamental asymmetries in brain function and behavior across vertebrate species, including humans and rodents^1^. Although individual hand or paw preferences are typically stable, they can be reshaped through experience or training, underscoring the plasticity of lateralized motor circuits^2^. However, the neural mechanisms that enable this behavioral switching remain unclear. Here we show that experience-driven shifts in paw preference in mice arise from asymmetric cholinergic modulation between the two hemispheres. Using a behavioral paradigm designed to induce paw switching, we found that a subset of animals developed a stable change in preference following training. Selective activation or inhibition of cholinergic neurons on one side of the brain was sufficient to bias this switching behavior. During lateralized movements, an endogenous imbalance in cholinergic activity emerges between the hemispheres, suggesting that interhemispheric cholinergic asymmetry enables motor flexibility. These findings identify a previously unrecognized mechanism by which cholinergic signaling dynamically regulates lateralized motor control, providing insight into how hemispheric imbalances may shape persistent individual motor preferences.

## Main

Motor skill lateralization reflects the functional asymmetry between the two hemispheres of the brain^3^. Motor preferences, such as handedness, represent a conserved behavioral expression of lateralization, observed across vertebrate species including humans and rodents^1^. In humans, hand preference is typically stable but can be modified through experience. Historically, left-handed individuals were often compelled to switch to their right hand, providing a unique context to study the plasticity of lateralized motor circuits. Although the behavioral and psychological consequences of such forced switching have been examined^4,5^, the neural mechanisms that enable shifts in hand preference remain poorly understood.

A potential contributor to this process is the acquisition of unilateral skills. The refinement of dexterous movements relies on the plasticity of the primary motor cortex (M1) in the contralateral hemisphere^6^, which in turn depends on cholinergic input from the nucleus basalis (NB) in the basal forebrain^7,8^. Notably, cholinergic projections from the NB are largely confined within each hemisphere^9,10^, suggesting that interhemispheric asymmetries in cholinergic tone may bias motor learning.

Similarly, rodents exhibit stable forelimb preference^2,11,12^, yet retain the capacity to switch paw use under specific conditions^2,13^. This behavioral flexibility makes them an ideal model for investigating the dynamics of lateralized motor control. Here, we established a paw-switching paradigm in mice to dissect the neural basis of experience-driven reorganization of lateralized motor output. We demonstrated that training induces a persistent shift in forelimb preference, driven by hemispheric asymmetry in cholinergic signaling.

### Sustained shift in paw preference following pellet-reach training

Although mice can adjust their paw use under biased task conditions^2,12^, it remains unclear whether this behavioral shift persists once the bias is removed. To address question, we developed a voluntary pellet-reaching task (Fig. 1a)^14,15^. The first two days served as the before-training phase, during which individual baseline paw preference was assessed, given the substantial variability in lateralized paw use among mice. Paw use was clearly lateralized, with 54.2% of the mice were classified as left-handed and 45.8% as right-handed (Fig. 1b), consistent with previous reports^11^. Fitting the data with a beta distribution (α = 0.010 and β = 0.024) confirmed a strongly U-shaped distribution of paw preference. This individual lateralization remained stable across sessions in nearly all mice (91.5%, 54 of 59 mice) (Fig. 1c).

**Fig. 1.**
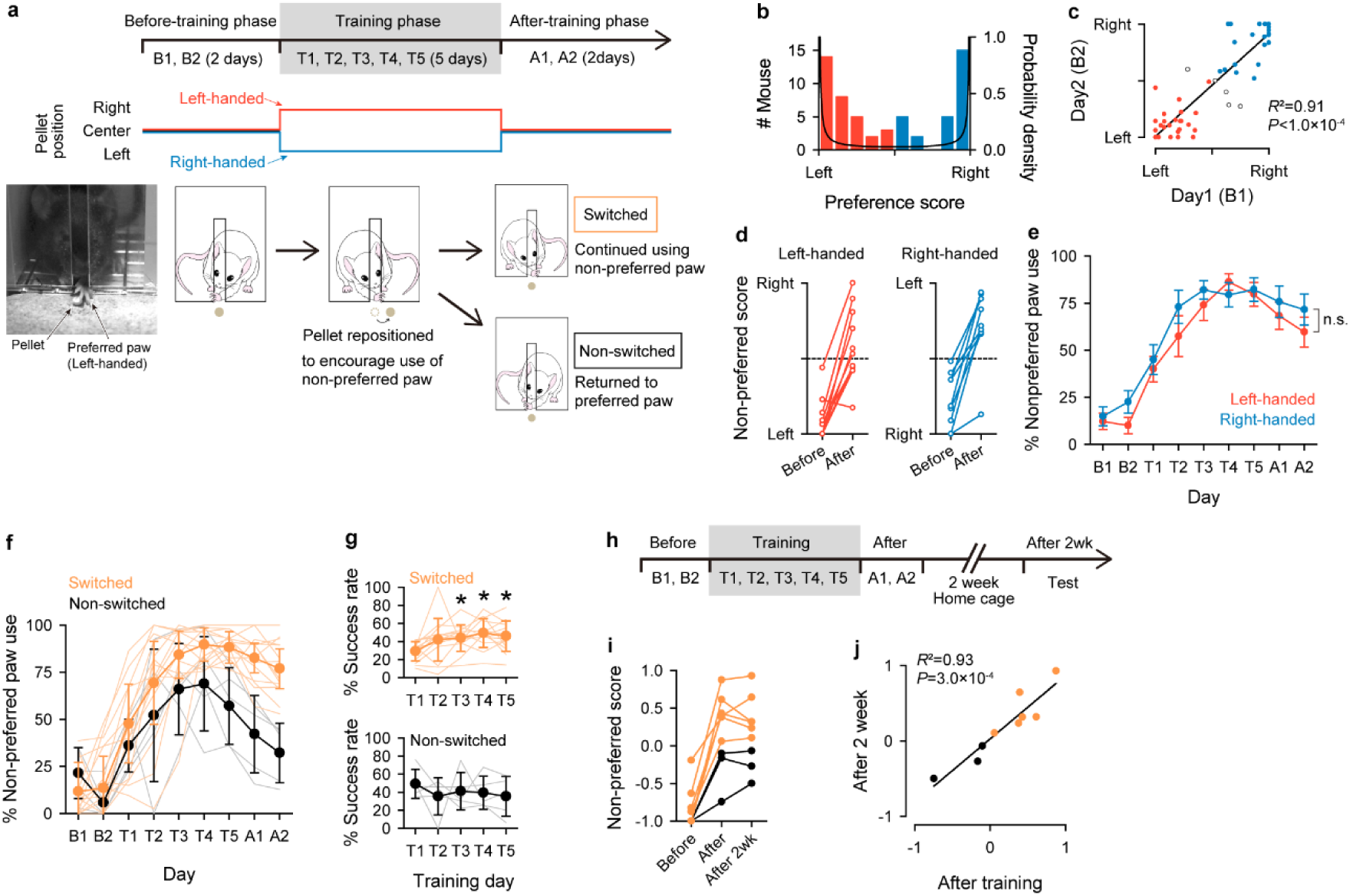
Paw switching through the training in mice. **a**, Behavioral paradigm of the paw-switching experiment. The preferred paw of each mouse was determined over a two-day period by a pellet-reaching task (before-training phase). During the five-day training phase, the pellet was laterally displaced so that mice could reach it only with their non-preferred paw (training phase). Finally, the pellet was returned to the center for a two-day period (after-training phase). **b**, Distribution of left- and right-handed mice. *n* = 59 mice. Beta distribution fitting yielded α = 0.010 and β = 0.024, indicating a strongly U-shaped distribution. **c,** Day 1 versus day 2 distribution of left- and right-handed mice showing stable individual lateralization, (*n* = 59 mice, linear regression). **d**, Non-preferred score, indicating the use rate of the non-preferred paw, increased in the after-training phase in both left- and right-handed mice (*n* = 10 left-handed and *n* = 9 right-handed mice). **e**, Transition in non-preferred paw use was comparable between left- and right-handed groups (*P* = 0.051, two-way ANOVA). **f**, Transition of non-preferred hand use in the switched and non-switched groups. **g**, Success rates during training. Switched group: *n* = 14, *P* = 1.7 × 10^-7^, non-switched group: *n* = 5, *P* = 0.80 (Tukey’s test following one-way ANOVA). **h**, Behavioral paradigm including an after-two-week test. **i**, Non-preferred score at three time points. **j**, Non-preferred scores remained stable two weeks after training (*n* = 9, *R*^2^ = 0.93, *P* = 3.0 × 10^-4^, linear regression).

During the training phase, pellet positions were shifted to prevent retrieval with the preferred paw, thereby prompting use of the non-preferred paw. Non-preferred paw use progressively increased over the five-day training period. After training, both left- and right-handed mice exhibited greater use of the non-preferred paw, although a subset continued to favor the original preference (Fig. 1d). This trend was consistent across both groups through all experimental phases (Fig. 1e). Because of this similarity, subsequent analyses pooled data from left- and right-handed animals. Based on the rate of non-preferred paw use during the after-training phase, mice were classified into switched or non-switched groups. Notably, these groups diverged early during the training phase (Fig. 1f). In the switched group, success rates improved over the training period, whereas the non-switched group showed little progress (Fig. 1g). These results suggest that enhancement of motor skills with the non-preferred paw can drive a stable switch in paw preference. Both newly acquired and pre-existing paw preferences persisted for at least two weeks following training (Fig. 1h–j).

### Unilateral NB lesion prevents paw switching

We next examined the role of the cholinergic system originating from the NB, which is essential for the improvement of forelimb skills^7,16^. This projection is predominantly ipsilateral^9,17,18^, a feature we confirmed using retrograde tracing with cholera toxin subunit B (CTb) and anterograde labeling with an adeno-associated virus (AAV) vector (Fig. 2a,b).

**Fig. 2.**
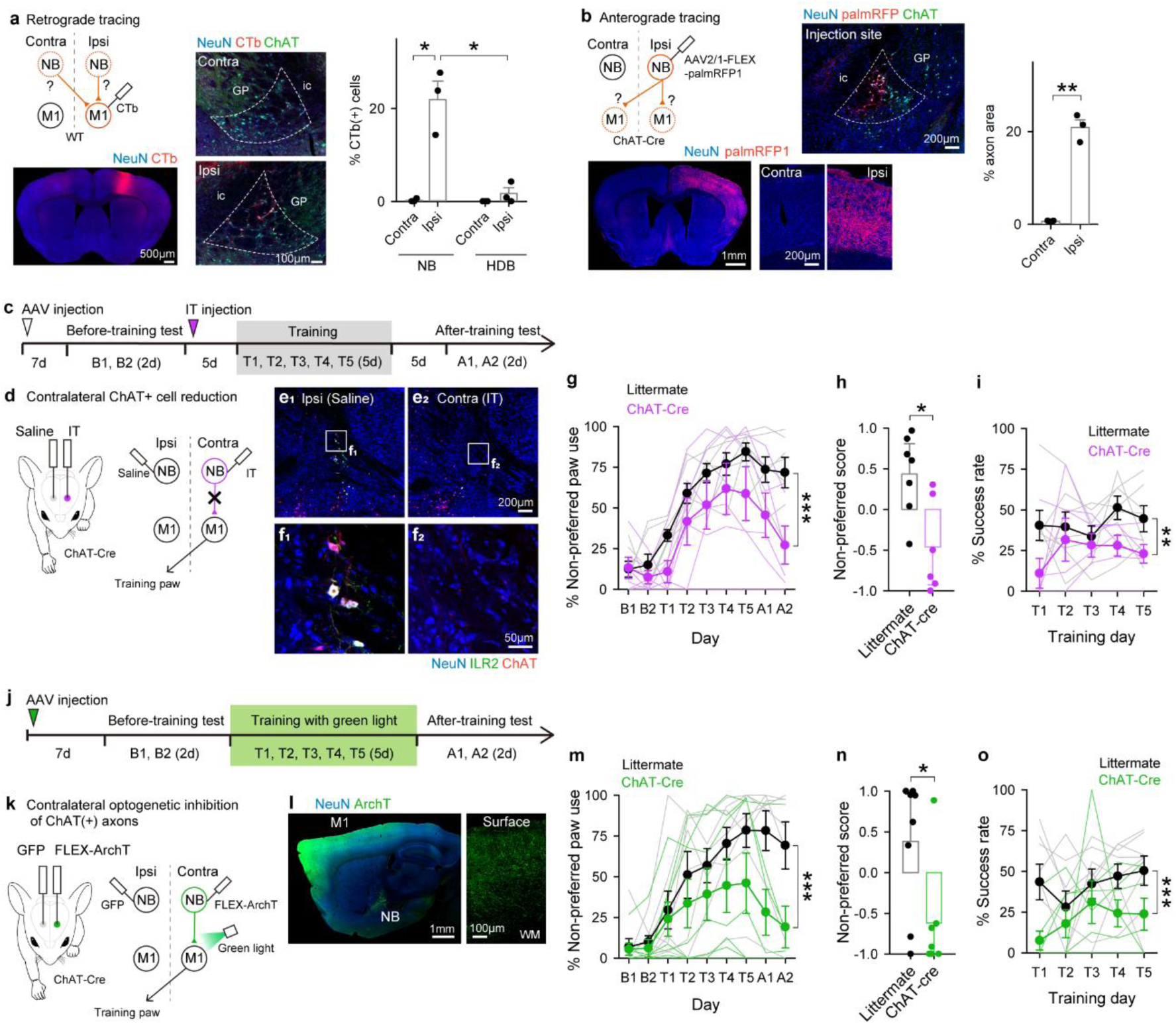
Paw switch requires contralateral cholinergic system. **a**, Retrograde tracing with CTb. CTb was unilaterally injected into the M1 forelimb area (left). CTb-positive cells were observed in the ipsilateral NB (right). NB is outlined by a white dashed line. GP, globus pallidus; ic, internal capsule. *n* = 3 mice, \**P* < 0.001 (Tukey’s test following two-way ANOVA). **b**, Anterograde tracing using an AAV vector. AAV2/1-FLEX-palmRFP1 was unilaterally injected into the NB of ChAT-Cre mice (top). Labeled axons projected ipsilaterally to the cortex (bottom; *n* = 3, \*\**P* = 7.1 × 10^-3^, paired *t*-test). **c**, Schematic of unilateral NB ablation. Immunotoxin (IT) was injected contralateral to the training paw, corresponding to non-preferred hand. **d**, Behavioral paradigm for NB ablation. **e–f**, Representative images of the NB ipsilateral (Ipsi) and contralateral (Contra) to the training paw. Cholinergic neurons were ablated in Contra NB. **g–i**, Behavioral effects of unilateral NB ablation. **g**, Transition in non-preferred paw use in ChAT-Cre mice and littermate controls (*n* = 7 littermates, *n* = 6 ChAT-cre mice, \*\*\**P* < 1.0 × 10^-3^, two-way ANOVA). **h**, Non-preferred score on day 9 (session A2; \**P* = 0.011, Student’s *t*-test). **i**, Success rate during training (\*\**P* = 2.0 × 10^-3^, two-way ANOVA). **j**, Schematic of unilateral optogenetic inhibition of NB-origin axons in M1. **k**, Behavioral diagram for optogenetic inhibition. **l**, Representative images showing ArchT expression in ipsilateral M1. WM, white matter. **m–o**, Behavioral outcomes of optogenetic inhibition. **m**, Transition in non-preferred hand use (*n* = 8 littermates, *n* = 7 ChAT-cre mice, \*\*\**P* < 1.0 × 10^-3^, two-way ANOVA). **n**, Non-preferred score on day 9 (session A2; \**P* = 0.025, Student’s *t*-test). **o**, Success rates during training (\*\*\**P* < 1.0 × 10^-3^, two-way ANOVA).

We hypothesized that paw switching is mediated by hemispheric asymmetry in cholinergic activity. To test this, we selectively ablated cholinergic neurons unilaterally in the NB contralateral to the training paw, using an immunotoxin system (anti-Tac(Fav)-PE83)^19,20^ in choline acetyltransferase (ChAT)-Cre transgenic mice (Fig. 2c–f). This unilateral cholinergic ablation significantly impaired paw switching compared with ChAT-Cre-negative littermates (Fig. 2g). The ablation group exhibited significantly lower non-preferred scores in the after-training phase (Fig. 2h)., and their success rates during training were also reduced (Fig. 2i). In contrast, ablation performed after training did not affect switching (Extended Data Fig. 1), consistent with previous findings^7^ that post-training cholinergic loss spares task performance. These findings indicate that cholinergic input is essential for enabling paw switching, and that its loss during the training phase disrupts motor learning performance.

Cholinergic axons from the NB innervate a wide range of brain areas within each hemisphere^21–23^. We therefore tested whether the NB-to-M1 pathway is required for paw switching. Optogenetic inhibition of cholinergic axon terminals in M1 during the training phase significantly impaired paw switching compared with ChAT-Cre-negative littermates (Fig. 2j–m). Non-preferred scores in the after-training phase were reduced (Fig. 2n), and success rates during training were similarly reduced (Fig. 2o). Together, these manipulations demonstrate that cholinergic input from the NB to M1 is indispensable for the acquisition of paw switching behavior.

Finally, we assessed whether unilateral manipulation of the NB in the opposite hemisphere affected paw switching. Immunotoxin injection ipsilateral to the training paw did not impair paw switching, but instead moderately facilitated it (Extended Data Fig.2), suggesting that ipsilateral ablation, unrelated to the training circuit, may influence motor preference via interhemispheric mechanisms.

### Task-evoked lateralization of cholinergic activity during paw use

To determine whether cholinergic activities in the two hemispheres are independent, we performed bilateral fiber photometry recordings. Cortical neurons were transduced with an AAV vector expressing gAch4h^24^, a genetically encoded fluorescent acetylcholine sensor, and signals were monitored bilaterally from the M1 as mice performed a voluntary pellet-reaching task (Fig. 3a). Contrary to expectations based on the predominantly unilateral organization of NB projections (Fig. 2a,b), cholinergic activity exhibited strong interhemispheric synchrony throughout the recording (Fig. 3b). When signals were aligned to the pellet-reaching movement with the preferred paw, both hemispheres showed concurrent increases inactivity, consistent with previous findings that acetylcholine levels rise during forelimb movement^25^. Notably, immediately after the reach, the hemisphere contralateral to the active paw displayed a transiently greater increase in signal intensity (Fig. 3b–d).

**Fig. 3.**
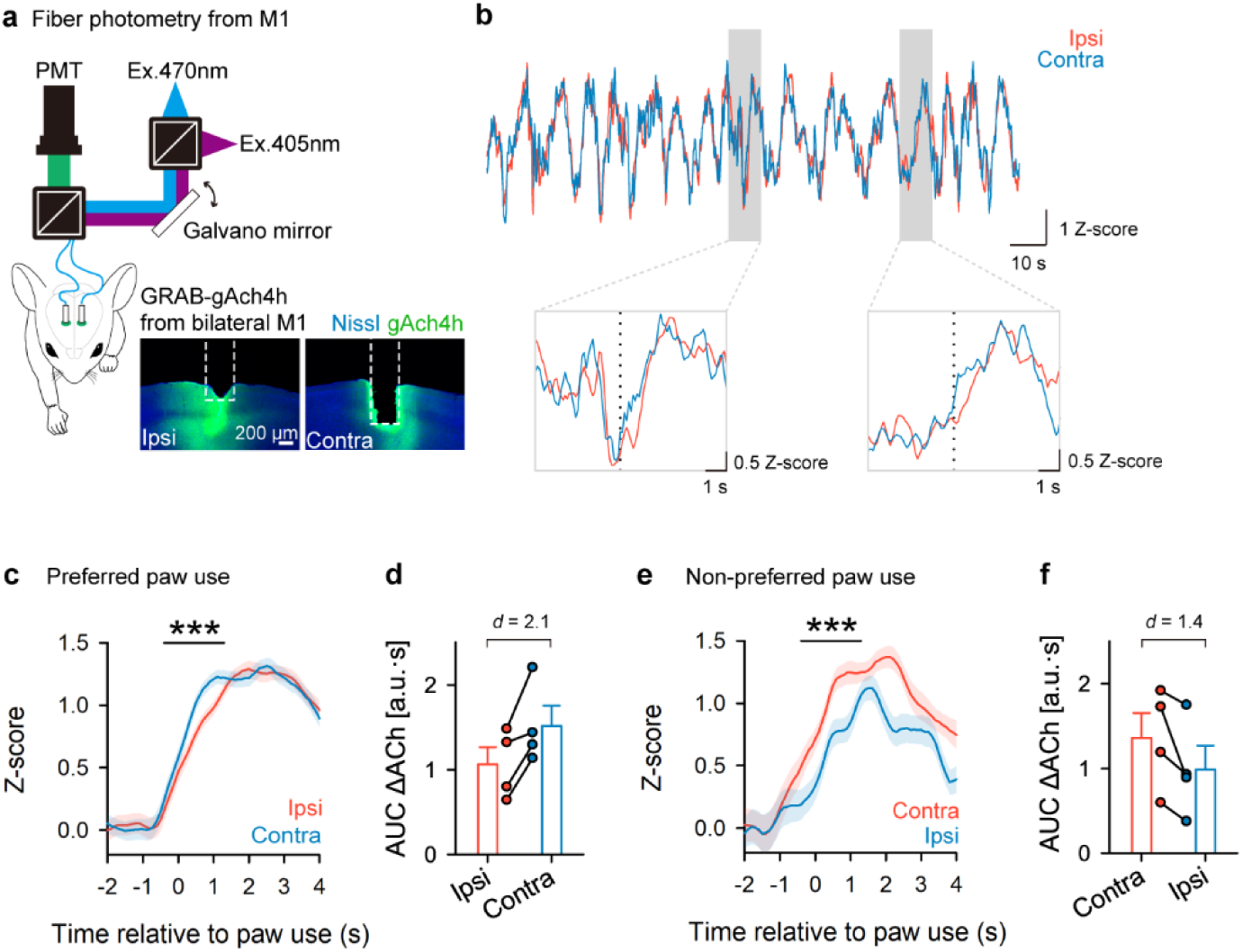
Task-evoked lateralization of cholinergic activity during paw use. **a**, Schematic of bilateral fiber photometry targeting M1. Representative images showing fiber tracks and GRAB-gAch4h expression in M1. **b**, Top: representative traces of gAch4h signals from ipsilateral (red) and contralateral (blue) M1. Bottom: enlarged view of shaded region. Dashed lines indicate paw-use onset. **c**, Averaged traces of gAch4h signals aligned to paw-use onset (*n* = 96 trials from 4 mice). Bar above indicates significant periods (\*\*\**P* = 2.0 ×10^-4^, cluster-based permutation test). **d**, Signal changes associated with paw use (*n* = 4 mice. Cohen’s *d* = 2.1). **e**, Same as (c), but for non-preferred paw use (*n* = 37 trials from 4 mice. \*\*\**P* = 8.0 ×10^-4^). **f**, Same as (d), but for non-preferred paw use (*n* = 4 mice. Cohen’s *d* = 1.4).

On the following day, when animals were trained to use the non-preferred paw, the hemisphere contralateral to the active paw, ipsilateral to the originally preferred paw, exhibited a stronger response than the opposite side (Fig. 3e,f). These results suggest that, while cholinergic activity in M1 is largely synchronized across hemispheres, it shows transient, task-locked lateralization contralateral to the engaged paw.

### Lateralized cholinergic balance impaired the paw switching

To experimentally induce asymmetry in cholinergic signaling, we unilaterally activated the NB using the DREADD system^26,27^. We first combined bilateral fiber photometry recordings from the M1 with chemogenetic activation to evaluate the effect of hM3Dq on cortical cholinergic dynamics (Extended Data Fig. 3a,b). Following intraperitoneal administration of deschloroclozapine (DCZ), the gAch4h signals increased predominantly in the M1 ipsilateral to the unilateral virus injection side relative to the contralateral hemisphere (Extended Data Fig. 3c).

We next expressed hM3Dq in the NB contralateral to the training paw and conducted a pellet-reaching task (Fig. 4a-c). DCZ was administered intraperitoneally each day during the five-day training phase. Chemogenetic activation of the contralateral NB did not alter paw-switching behavior (Fig. 4d,e), although activated mice exhibited higher success rates than controls (Fig. 4f).

**Fig. 4.**
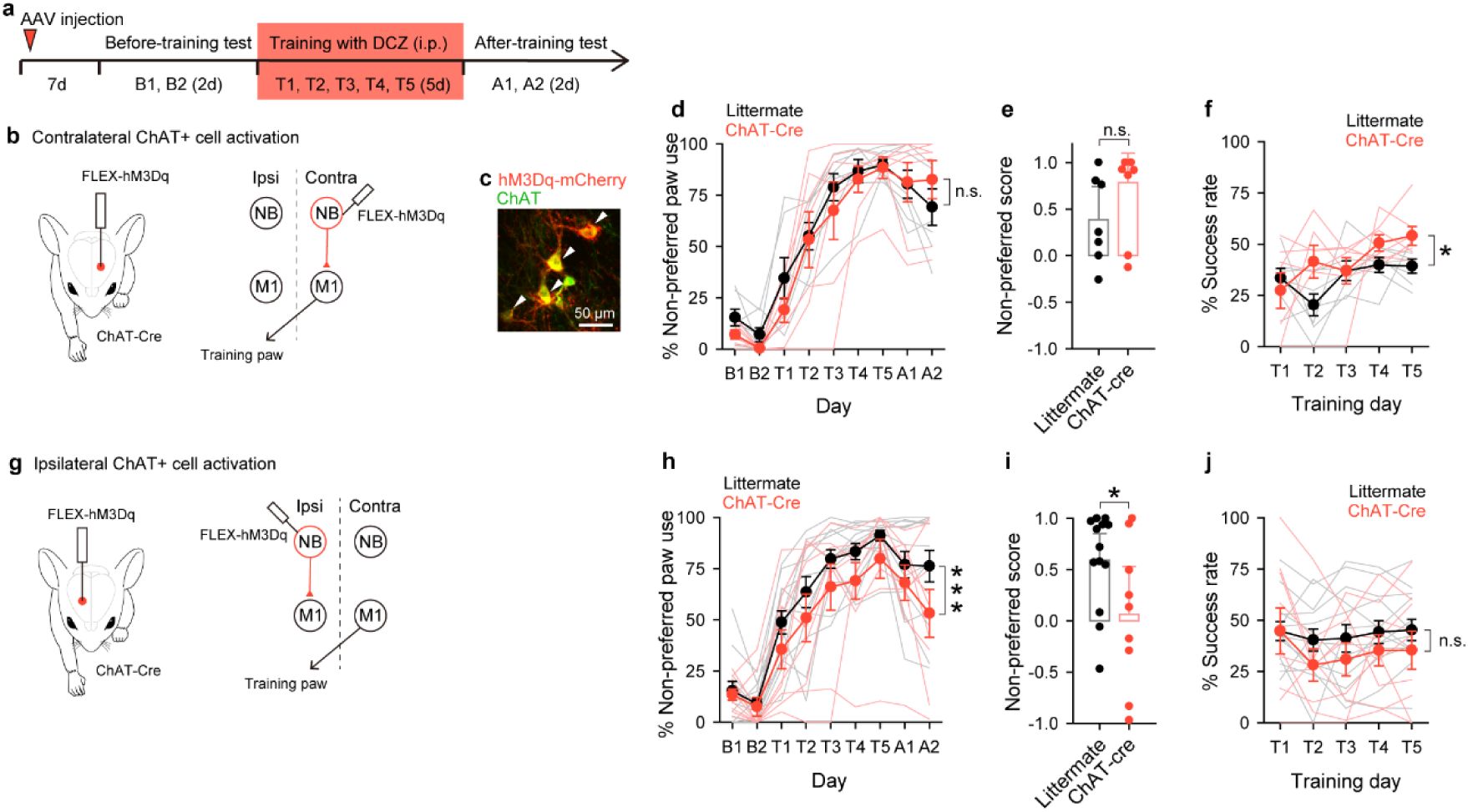
Interhemispheric balance of cholinergic activity. **a**, Schematic of unilateral NB activation. **b**, Behavioral paradigm for contralateral NB activation. **c**, Representative image of hM3Dq-expressing cells in NB. **d–f,** Behavioral outcomes of contralateral NB activation. **d,** Transition in non-preferred paw use (*n* = 7 littermates, *n* = 6 ChAT-cre mice, *P* = 0.29, two-way ANOVA). **e,** Non-preferred score on day9 (session A2; *P* = 0.13, Student’s *t*-test). **f,** Success rates during training (\**P* = 0.027, two-way ANOVA). **g,** Behavioral paradigm for ipsilateral NB activation. **h–j,** Behavioral outcomes of ipsilateral NB activation. **h**, Transition in non-preferred hand use (*n* = 13 littermates, *n* = 9 ChAT-cre mice, \*\*\**P* < 1.0 × 10^-3^, two-way ANOVA). **i**, Non-preferred score on day 9 (session A2; \**P* = 0.045, Student’s *t*-test). **j**, Success rates during training (*P* = 0.068, two-way ANOVA).

In contrast, activation of the NB ipsilateral to the training paw suppressed paw switching (Fig. 4g–i), without affecting overall task performance (Fig. 4j). This effect parallels that of ipsilateral immunotoxin ablation (Extended Data Fig. 2), suggesting that unbalanced cholinergic activation across hemispheres biases motor preference through interhemispheric interactions. Collectively, these findings indicate that improvements in task performance alone cannot account for the induction of paw switching; rather, an interhemispheric cholinergic dominance is required.

## Discussion

In this study, we investigated the role of interhemispheric cholinergic modulation in paw preference switching. We established a behavioral paradigm in which mice exhibited training-induced changes in paw preference, revealing that this process requires cholinergic input from the NB, which exerts hemispheric-dominant control over M1. Although acetylcholine dynamics in M1 were largely synchronized bilaterally, a transient lateralized enhancement contralateral to the reaching paw emerged during task execution. Notably, manipulation of the hemisphere not directly engaged in the trained paw movement also influenced switching, suggesting that lateralized motor behavior is governed by the interhemispheric balance of cholinergic activity.

Although forelimb preference in rodents is well-documented, how it can be modified through training remains unclear. Using our developed behavioral paradigm, we demonstrated that repeated training induces a robust and persistent switch in paw use, with the newly acquired preference maintained for at least two weeks (Fig. 1h–j). Consistent with the individually determined nature of paw preference, the paradigm revealed substantial inter-individual variability. Many mice adopted and maintained the trained paw, whereas others either retained their original preference or reverted after transient switching (Fig. 1f). Such heterogeneity likely reflects intrinsic differences in the flexibility of lateralized motor circuits.

Unilateral skill enhancement is required for paw-switching, a process depends on cholinergic signaling^7,8,25^, and hemispheric dominance of projections from the NB^9,10,18^. Consistent with this, unilateral disruption of cholinergic input prevented paw switching (Fig. 2). However, motor improvement alone was insufficient to induce switching, as shown in Fig. 4. Chemogenetic activation of cholinergic input contralateral to the training paw increased success rates but did not affect switching (Fig. 4b–f), whereas ipsilateral activation suppressed switching without impairing performance (Fig. 4g–j). These findings suggest that cholinergic signaling contributes not only to the facilitation of motor learning but also to circuit-level processes that reorganize lateralized motor output.

Basal forebrain cholinergic activity has been implicated in a range of functions, including arousal, attention, and learning^7,28–30^. Given this diversity, the gAch4h signals recorded here may reflect multiple components beyond motor-specific modulation, potentially accounting for the high degree of interhemispheric synchrony observed (Fig. 3b). Nevertheless, anatomical evidence indicates that cholinergic projections from the NB to M1 are largely confined within hemispheres (Fig. 2a,b)^9,10^, and that these M1-projecting cholinergic neurons receive input exclusively from the striatum, which itself exhibits lateralized connectivity^22^. Therefore, the task-locked contralateral enhancements observed during single-paw reaching may represent a motor-related component of this broader cholinergic activity (Fig. 3c–f). Repeated experience may initially amplify subtle cholinergic asymmetries, leading to the stabilization of lateralized motor output over time.

Finally, our findings suggest that interhemispheric asymmetry in cholinergic activity contributes to the regulation of motor lateralization. Although cholinergic projections from the NB to M1 are predominantly unilateral (Fig. 2a,b), non-cholinergic neurons within the NB have been reported to form reciprocal connections across hemispheres^31,32^, and cortical areas are extensively interconnected through commissural fibers^33–35^. These interhemispheric connections may permit competitive interactions between the hemispheres, thereby facilitating lateralized motor execution^36^. Therefore, even subtle imbalances in cholinergic activity during unilateral training may gradually bias motor output and consolidate asymmetric motor behaviors at the individual level.

## Methods

### Animals

All animal experiments were conducted in accordance with the National Institutes of Health Guide for the Care and Use of Laboratory Animals and were approved by the Committees for Animal Care and Use and Recombinant DNA Study at Juntendo University (2024231, 2024232, 2024213, and 2024213). C57BL/6J mice (male and female; 6–10 weeks old) were obtained from Nihon SLC. ChAT-Cre(BAC) mice (male and female; 6–16 weeks old) were obtained from The Jackson Laboratory and maintained by continuous breeding. Animals were housed under a 12-h light/dark cycle with ad libitum access to food and water. All procedures were designed to minimize animal suffering and reduce the number of animals used. This study was conducted in compliance with the ARRIVE guidelines.

### Pellet reaching task

Mice were food-restricted to maintain 85–90% of their free-feeding weight before the start of training. The behavioral chamber was an acrylic box (115 mm tall × 55 mm deep × 38 mm wide) with a transparent front wall and opaque walls on the remaining three sides. A vertical slit (5 mm wide × 57.5 mm high) was located at the center of the front wall. A food pellet was positioned on a cork plate, 5–7 mm from the slit^14,15^. During training, the pellet was placed 5 mm to the left or right of the slit center, according to each mouse’s initial paw preference. On the day prior to behavioral testing, each mouse was habituated to the chamber for 10 minutes.

The behavioral procedure consisted of three phases. In the before-training phase (2 days), the preferred paw of each mouse was determined using a pellet-reaching task. During the training phase (5 days), the pellet was repositioned laterally so that the mouse could reach it only with its non-preferred paw. In the after-training phase (2 days), the pellet was returned to the center position in front of the slit. Each daily behavioral session lasted 10 minutes. For the immunotoxin (IT) experiments, an additional 5-day interval was introduced between each phase to allow for cellular reduction.

Behavioral performance for one or two mice was recorded frontally using a monochrome camera (BU205M; Toshiba Teli) at 100 Hz. The number of reaching attempts with the right and left paws, as well as the success rate, were manually assessed. In the before-training phase, the paw with the greater number of reaching attempts was defined as the preferred paw. Mice that switched their preferred paw between sessions B1 and B2 were excluded from subsequent analyses (from Fig. 1d onward).

The percentage of non-preferred paw use was calculated as:

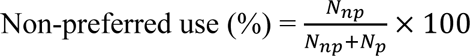

The non-preferred score was calculated as:

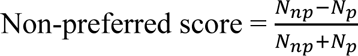

where N_np_ and N_p_ represent the number of reaching attempts with the non-preferred and preferred paw, respectively. The success rate was calculated as the number of successful pellet retrievals divided by the total number of attempts. As shown Fig. 1, mice were categorized as switched when their non-preferred score in session A2 was positive and as non-switched when it was negative.

### Production and purification of AAV vectors

AAV vector particles were produced and purified as described previously^37–39^. HEK293T cells were incubated in 10 CELLSTAR^®^ cell culture dishes (145 mm; #639160, Greiner Bio-One). Each genomic plasmid and two helper plasmids, pBSIISK-R2C1^40^ and pHelper (28060929; Stratagene) were co-transfected into HEK293T cells using polyethylenimine (23966; Polysciences). Virus particles were harvested from both the cell lysate and the supernatant, followed by ultracentrifugation with OptiPrep (Axis-Shield). The purified solution was ultrafiltrated and concentrated in Dulbecco’s phosphate-buffered saline (PBS; 14249-95; Nacalai Tesque) containing 0.001% Pluronic F-68 (24040032; Thermo Fisher Scientific) using an Amicon Ultra-15 (NMWL 30K; Merck Millipore). Following the ultrafiltration, each virus solution was concentrated to a final volume of 100 µL. The virus titer (gc/mL) was quantified using qPCR with Fast SYBR^®^ Green Master Mix (#4385612, Applied Biosystems). The final titers were 8.4 × 10^11^ gc/mL for AAV2/1-CAGGS-FLEX-[IL2R-EGFP]-3’USS^19,20^, 1.4 × 10^15^ gc/mL for AAV2/1-CAG-FLEX-[ArchT-EGFP]^41^, 8.4 × 10^13^ gc/mL for AAV2/1-CAG-FLEX-[hM3Dq-mCherry]^27^, 7.7 × 10^14^ gc/mL for AAV2/1-hSyn-gAch4h^24^, and 3.6 × 10^14^ gc/mL for AAV2/1-SynTetOff-FLEX-[palmRFP1]^40^. The virus solutions were stored in aliquots at -80°C until use.

### Stereotaxic surgical procedures

Mice were deeply anesthetized by intraperitoneal injection of a mixture of medetomidine (0.3 mg/kg; Nippon Zenyaku Kogyo), midazolam (4 mg/kg; Sandoz), and butorphanol (5 mg/kg; Meiji Seika Pharma) and placed in a stereotaxic apparatus. We injected 0.2 µL of the solution into the NB (0.5 mm posterior to the bregma, ±1.8 mm lateral to the midline, 4.5 mm ventral to the brain surface) and/or the M1 forelimb area (0.5 mm anterior to the bregma, ±1.7 mm lateral to the midline, 0.8 mm ventral to the brain surface) by pressure through a glass micropipette attached to Picospritzer III (Parker Hannifin).

For retrograde tracing, cholera toxin subunit b (CTb; #103C, List Biological Laboratories) was used at a concentration of 2% (w/v).

For IT-mediated cell reduction^19,20,42^, an IT solution (5 µg/µl anti-Tac(Fav)-PE38 in PBS containing 0.1% bovine serum albumin) was injected into the NB of animals previously injected with AAV2/1-CAGGS-FLEX-[IL2R-EGFP]-3’USS, after a delay of more than one week.

For optogenetic manipulation, a 1-mm-diameter cranial window was opened over the M1 injection site. A green light-emitting diode (LED) device of TeleOpto system (TeleLDP-c-i; Bio Research Center) was implanted and secured to the skull with dental cement. A 2-g dummy receiver (TeleDummy, Bio Research Center) was attached to the LED device.

For fiber photometry, optic fiber cannulas (400 µm diameter, 0.5 NA, R-FOC-BL400C-50NA; RWD) were bilaterally implanted into the M1 forelimb area following viral injection. Fiber ferrules were fixed to the skull with a quick-setting adhesive (LOCTITE 454; Henkel) and coated with a black surfacer (GS-03; Gianotes).

After surgery, mice received an intraperitoneal injection of atipamezole (Antisedan; 1.5 mg/kg; Orion) and recovered from anesthesia within approximately 15 minutes.

### Optogenetic manipulation

The TeleOpt system (Bio Research Center) was used for optogenetic manipulations. Before the behavioral experiments, the wireless receiver (TeleR-2-C; Bio Research Center) was replaced with a dummy and connected to the LED device. Green light was continuously delivered throughout the 10-minute training sessions. After the behavioral experiment, the wireless receiver was again replaced with the dummy unit.

### Fiber photometry

Fiber photometry recordings were performed using bifurcated fiber-optic bundles (400 µm diameter, 0.39 NA; BFYL4LF01, Thorlabs) attached to the optic fiber ferrules and covered with black sleeves (Lymyth). Recordings were conducted using a Thorlabs fiber photometry system, which delivered excitation light (470 nm for the active signal and 405 nm for the isosbestic reference) through the patch cord as interleaved LED pulses at 40 Hz. The two optical paths alternated at 20 Hz using a galvanometric mirror (GVS001, Thorlabs), resulting in an acquisition rate of 10 Hz per channel. Light was reflected through a dichroic mirror (DMLP490R, Thorlabs) and detected using a photomultiplier tube (PMT1001/M, Thorlabs). Excitation power was adjusted at ∼50-90 µW at the patch cord tip. Signals were acquired with a data acquisition interface (USB-6003, National Instruments) and custom-written LabVIEW software (National Instruments). Fiber photometry recordings began immediately after the start of behavioral video acquisition. Synchronization between the two systems was achieved by capturing a 40-Hz-blinking infrared LED within the video frame.

During the recordings, mice performed a pellet-reaching task for 10 minutes. Photometry data were collected across two sessions with the preferred paw, during which the pellet was placed at the center of the slit. On a separate day, recordings of the non-preferred paws were obtained by laterally shifting the pellet position.

The photometry signals analyzing using custom MATLAB scripts (MathWorks). Raw fluorescence traces from the 470 nm and 405 nm channels were median-filtered; the 470-nm active channel was normalized by subtracting the 405-nm reference signal, low-pass filtered, and converted to z-scores. Processed signals were time-locked to successful paw-reaching events for event-related analysis.

### DREADD manipulation

Deschloroclozapine (HY-42110, MedChemExpress) was dissolved in saline containing 0.1% dimethyl sulfoxide (DMSO) to a final concentration of 2 ug/mL and intraperitoneally administrated at a 0.1 mL per mouse, 10–20 minutes before each behavioral session during the training phase^26^. Animals received unilateral injections of AAV2/1-CAG-FLEX-[hM3Dq-mCherry] into NB 1–2 weeks before the start of the behavioral testing.

### Histology

Mice were deeply anesthetized by intraperitoneal injection of sodium pentobarbital (200 mg/kg; Somnopentyl, Kyoritsu Seiyaku) and perfused transcardially with 20 mL of PBS (pH 7.4, 20–25°C), followed by perfusion for 3 minutes with 4% paraformaldehyde (1.04005.1000; Millipore) in 0.1 M phosphate buffer (PB; pH 7.4, 20–25°C). The brains were removed and post fixed overnight at 4°C in the same fixative. After cryoprotection in 30% sucrose in 0.1 M PB, the brains were cut into 40-µm-thick parasagittal or coronal sections on a freezing microtome (REM-710; Tamato Kohki Industrial). Sections were collected in six bottles containing 0.02% sodium azide in PBS and stored at 4°C until use for free-floating immunostaining.

All following incubations were performed at 20–25°C and followed by rinsing twice with PBS containing 0.3% (v/v) Triton X-100 (PBS-X) for 10 minutes. Free-floating sections (40-µm thick) were incubated for 16 hours with 1:1000-diluted goat polyclonal antibody against ChAT (AB144P, Millipore), 1:1000-diluted mouse monoclonal antibody against NeuN (MAB377, Millipore), 1:500-diluted rabbit polyclonal antibody against ChAT (AB143, Millipore), 1:1000-diluted goat polyclonal antibody against CTb, 1:1000-diluted affinity-purified rabbit antibody against mRFP^43^, or 1:100-diluted chicken polyclonal antibody against GFP (GFP-1020, Aves Labs) in PBS-X containing 1% (v/v) donkey serum (S30-100ML; Merck) and 0.12% (w/v) λ-carrageenan (035-09693; Wako Chemicals) (PBS-XCD). The sections and slices were incubated for 4 h with 5 μg/mL Alexa Fluor (AF) 647-conjugated donkey anti-goat IgG (ab150131; Abcam), AF405-conjugated donkey anti-mouse IgG (ab175658; Abcam), AF568-conjugated donkey anti-goat IgG (A11057; Thermo Fisher Scientific), AF647-conjugated donkey anti-rabbit IgG (A10042; Thermo Fisher Scientific), AF647-conjugated donkey anti-chicken IgG (A21469; Thermo Fisher Scientific), or AF568-conjugated donkey anti-rabbit IgG (A10042; Thermo Fisher Scientific) in PBS-XCD, followed by incubation for 30 min with 1:300-diluted NeuroTrace 435/455 Blue Fluorescent Nissl stain (N21479; Thermo Fisher Scientific) in PBS-X.

The sections were imaged using a confocal laser scanning microscope (TCS SP8; Leica Microsystems) equipped with 10× (HCX PL APO 10x/0.40 CS, NA = 0.40; Leica Microsystems), 16× multi-immersion (HC FLUOTAR 16x/0.60 IMM CORR VISIR, NA = 0.60; Leica Microsystems), and 25× water-immersion (HC FLUOTAR L 25x/0.95 W VISIR, NA = 0.95; Leica Microsystems) objective lenses. NeuroTrace Blue, EGFP or AF488, AF568, mRFP1, or mCherry, and AF647 were excited using 405-, 488-, 552-, and 638-nm laser lines, respectively. Emission signals were collected through 410–503, 495–550, 570–650, and 660–750-nm emission prism windows, and detected in photon-counting mode using hybrid detector (HyD; Leica Microsystems).

### Statistics and reproducibility

Data are presented as mean ± standard error of the mean (SEM). The exact values of *n* are provided in the corresponding figure legends. Statistical analyses were conducted using MATLAB (MathWorks) and SigmaPlot (HULINKS). Two-way analysis of variance (ANOVA) was used to compare daily changes between groups. One-way ANOVA followed by Tukey’s post hoc test was applied for within-group changes across days (Fig. 1g). Student’s *t*-tests were used for independent pairwise group comparisons. Paired *t*-test was used for pairwise group comparisons (Extended Data Fig. 3c). Linear regression was used to compare distributions across two days (Fig. 1c,j). Cluster-based permutation tests were used to evaluate temporal differences in bilateral activity (Fig. 3c,e). Cohen’s d was calculated to estimate the effect sizes of differences in bilateral activity (Fig. 3d,f). All tests were two-sided, and statistical significance was set at *P* < 0.05.

## Acknowledgments

We thank T. Imura (Juntendo University) for technical assistance. This study was supported in part by KAKENHI (JP20K16112 and JP22K15230 to K.O.; JP21H02592, JP23K20044 and JP25K1856 to H.H.) from the Japan Society for the Promotion of Science (JSPS). This study was also supported by the Japan Agency for Medical Research and Development (AMED) (JP23wm0625001 and JP24wm0625103 to H.H.), PRESTO from the Japan Science and Technology Agency (JST) (JPMJPR23S3 to K.O.), Moonshot R&D from the JST (JPMJMS2024 to H.H.), Fusion Oriented Research for disruptive Science and Technology (FOREST) from JST (JPMJFR204D to H.H.), Grants-in-Aid from the Research Institute for Diseases of Old Age at the Juntendo University School of Medicine (X2208, X2311, and X2411 to K.O.), and the Private School Branding Project.

## Author contributions

Conceptualization, K.O. and H.H.; Methodology, K.O., Y.R.T., and H.H.; Investigation, K.O.; Resources, S.K., S.X., G.L., K.K., Y.L., and H.H.; Visualization, K.O.; Writing - Original Draft, K.O. and H.H.; Writing - Review & Editing, K.O., Y.R.T., S.K., S.X., G.L., K.K., Y.L., M.K., and H.H.; Project Administration, K.O. and H.H; Funding Acquisition, K.O. and H.H.

## Data and code availability

All data and custom scripts used in this study are available from the corresponding author on reasonable request.

## References

1. Stancher, G., Sovrano, V. A. & Vallortigara, G. Motor asymmetries in fishes, amphibians, and reptiles. Prog. Brain Res. 238, 33–56 (2018).

2. Collins, R. When left-handed mice live in right-handed worlds. Science 187, 181–184 (1975).

3. Güntürkün, O., Ströckens, F. & Ocklenburg, S. Brain lateralization: A comparative perspective. Physiol. Rev. 100, 1019–1063 (2020).

4. Klöppel, S., Mangin, J. F., Vongerichten, A., Frackowiak, R. S. J. & Siebner, H. R. Nurture versus nature: Long-term impact of forced right-handedness on structure of pericentral cortex and basal ganglia. Journal of Neuroscience 30, 3271–3275 (2010).

5. Sun, Z. Y. et al. The effect of handedness on the shape of the central sulcus. Neuroimage 60, 332–339 (2012).

6. Kawai, R. et al. Motor Cortex Is Required for Learning but Not for Executing a Motor Skill. Neuron 86, 800–812 (2015).

7. Conner, J. M., Culberson, A., Packowski, C., Chiba, A. A. & Tuszynski, M. H. Lesions of the basal forebrain cholinergic system impair task acquisition and abolish cortical plasticity associated with motor skill learning. Neuron 38, 819–829 (2003).

8. Conner, J. M., Kulczycki, M. & Tuszynski, M. H. Unique contributions of distinct cholinergic projections to motor cortical plasticity and learning. Cereb. Cortex 20, 2739–2748 (2010).

9. Kim, J. H. et al. Selectivity of neuromodulatory projections from the basal forebrain and locus ceruleus to primary sensory cortices. Journal of Neuroscience 36, 5314–5327 (2016).

10. Wu, H., Williams, J. & Nathans, J. Complete morphologies of basal forebrain cholinergic neurons in the mouse. Elife 2014, 1–17 (2014).

11. Manns, M., Basbasse, Y. E., Freund, N. & Ocklenburg, S. Paw preferences in mice and rats: Meta-analysis. Neurosci. Biobehav. Rev. 127, 593–606 (2021).

12. Tang, A. C. & Verstynen, T. Early life environment modulates “handedness” in rats. Behav. Brain Res. 131, 1–7 (2002).

13. Stieger, B., Palme, R., Kaiser, S., Sachser, N. & Richter, S. H. When left is right: The effects of paw preference training on behaviour in mice. Behav. Brain Res. 430, 113929 (2022).

14. Xu, T. et al. Rapid formation and selective stabilization of synapses for enduring motor memories. Nature 462, 915–919 (2009).

15. Sohn, J. et al. Presynaptic supervision of cortical spine dynamics in motor learning. Science Advances 8, (2022).

16. Li, Y. & Hollis, E. Basal Forebrain Cholinergic Neurons Selectively Drive Coordinated Motor Learning in Mice. J. Neurosci. 41, 10148–10160 (2021).

17. Casamenti, F., Di Patre, P. L., Bartolini, L. & Pepeu, G. Unilateral and bilateral nucleus basalis lesions: differences in neurochemical and behavioural recovery. Neuroscience 24, 209–215 (1988).

18. Buzsaki, G. et al. Nucleus basalis and thalamic control of neocortical activity in the freely moving rat. J. Neurosci. 8, 4007–4026 (1988).

19. Okada, K. et al. Enhanced flexibility of place discrimination learning by targeting striatal cholinergic interneurons. Nat. Commun. 5, (2014).

20. Okada, K., Nishizawa, K., Kobayashi, T., Sakata, S. & Kobayashi, K. Distinct roles of basal forebrain cholinergic neurons in spatial and object recognition memory. Sci. Rep. 5, 13158 (2015).

21. Chen, Z.-Y. et al. Whole-brain neural connectivity to cholinergic neurons in the nucleus basalis of Meynert. J. Neurochem. (2023) doi:10.1111/jnc.15873.

22. Gielow, M. R. & Zaborszky, L. The Input-Output Relationship of the Cholinergic Basal Forebrain. Cell Rep. 18, 1817–1830 (2017).

23. Li, X. et al. Generation of a whole-brain atlas for the cholinergic system and mesoscopic projectome analysis of basal forebrain cholinergic neurons. Proc. Natl. Acad. Sci. U. S. A. 115, 415–420 (2017).

24. Costa, K. M. et al. Dopamine and acetylcholine correlations in the nucleus accumbens depend on behavioral task states. Curr. Biol. 35, 1400–1407.e3 (2025).

25. Ren, C. et al. Global and subtype-specific modulation of cortical inhibitory neurons by acetylcholine during motor learning. Neuron 4 (2022).

26. Nagai, Y. et al. Deschloroclozapine, a potent and selective chemogenetic actuator enables rapid neuronal and behavioral modulations in mice and monkeys. Nat. Neurosci. 23, 1157–1167 (2020).

27. Krashes, M. J. et al. Rapid, reversible activation of AgRP neurons drives feeding behavior in mice. J. Clin. Invest. 121, 1424–1428 (2011).

28. Hangya, B., Ranade, S. P., Lorenc, M. & Kepecs, A. Central Cholinergic Neurons Are Rapidly Recruited by Reinforcement Feedback. Cell 162, 1155–1168 (2015).

29. Laszlovszky, T. et al. Distinct synchronization, cortical coupling and behavioral function of two basal forebrain cholinergic neuron types. Nat. Neurosci. 23, 992–1003 (2020).

30. Xu, M. et al. Basal forebrain circuit for sleep-wake control. Nat. Neurosci. 18, 1641–1647 (2015).

31. Semba, K., Reiner, P. B., McGeer, E. G. & Fibiger, H. C. Non-cholinergic basal forebrain neurons project to the contralateral basal forebrain in the rat. Neurosci. Lett. 84, 23–28 (1988).

32. Do, J. P. et al. Cell type-specific long-range connections of basal forebrain circuit. Elife 5, 1–18 (2016).

33. Oka, Y. et al. Interstitial Axon Collaterals of Callosal Neurons Form Association Projections from the Primary Somatosensory to Motor Cortex in Mice. Cereb. Cortex 31, 5225–5238 (2021).

34. Ocklenburg, S. & Guo, Z. V. Cross-hemispheric communication: Insights on lateralized brain functions. Neuron (2024) doi:10.1016/j.neuron.2024.02.010.

35. Handa, T., Zhang, Q. & Aizawa, H. Cholinergic modulation of interhemispheric inhibition in the mouse motor cortex. Cereb. Cortex 34, (2024).

36. Fenk, L. A., Riquelme, J. L. & Laurent, G. Interhemispheric competition during sleep. Nature (2023) doi:10.1038/s41586-023-05827-w.

## Reference

37. Takahashi, M., Ishida, Y., Kataoka, N., Nakamura, K. & Hioki, H. Efficient Labeling of Neurons and Identification of Postsynaptic Sites Using Adeno-Associated Virus Vector. vol. 169 (2021).

38. Furuta, T., et al. Multi-scale light microscopy/electron microscopy neuronal imaging from brain to synapse with a tissue clearing method, ScaleSF. iScience 25, 103601 (2022).

39. Okamoto, K., et al. Specific AAV2/PHP.eB-mediated gene transduction of CA2 pyramidal cells via injection into the lateral ventricle. Sci. Rep. 13, 323 (2023).

40. Sohn, J., et al. A single vector platform for High-Level gene transduction of central neurons: Adeno-associated virus vector equipped with the Tet-Off system. PLoS One 12, e0169611 (2017).

41. Han, X., et al. A high-light sensitivity optical neural silencer: development and application to optogenetic control of non-human primate cortex. Front. Syst. Neurosci. 5, 18 (2011).

42. Kreitman, R. J., Bailon, P., Chaudhary, V. K., FitzGerald, D. J. & Pastan, I. Recombinant immunotoxins containing anti-Tac(Fv) and derivatives of Pseudomonas exotoxin produce complete regression in mice of an interleukin-2 receptor-expressing human carcinoma. Blood 83, 426–434 (1994).

43. Hioki, H., et al. Vesicular glutamate transporter 3-expressing nonserotonergic projection neurons constitute a subregion in the rat midbrain raphe nuclei. J. Comp. Neurol. 518, 668–686 (2010).

