## Supplemental figures for "Paw switching with lateralized cholinergic modulation"

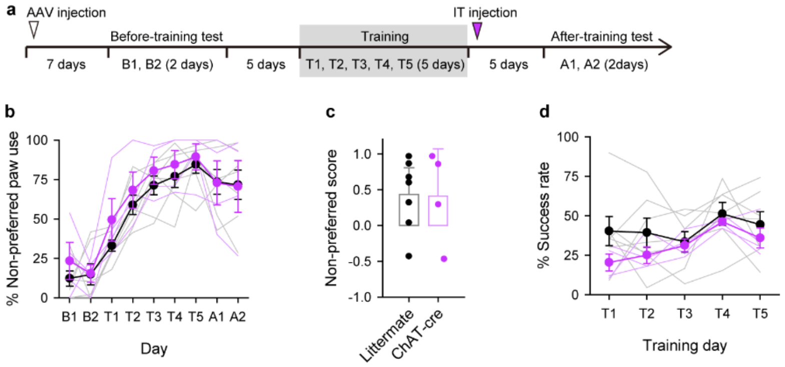


**Extended Data Fig. 1 | Contralateral NB ablation after training does not affect paw switching.**

**a**, Behavioral paradigm for NB ablation. **b**, Transition in non-preferred paw use in ChAT-Cre mice and littermate controls. Littermate controls were from the same cohort used in Figure 2**c–i** (*n* = 7 littermates, 4 ChAT-cre mice, *P* = 0.12, two-way ANOVA). **c**, Non-preferred score on day 9 (session A2; *P* = 0.95, Student's *t*-test). **d**, Success rates during training (*P* = 0.060, two-way ANOVA).

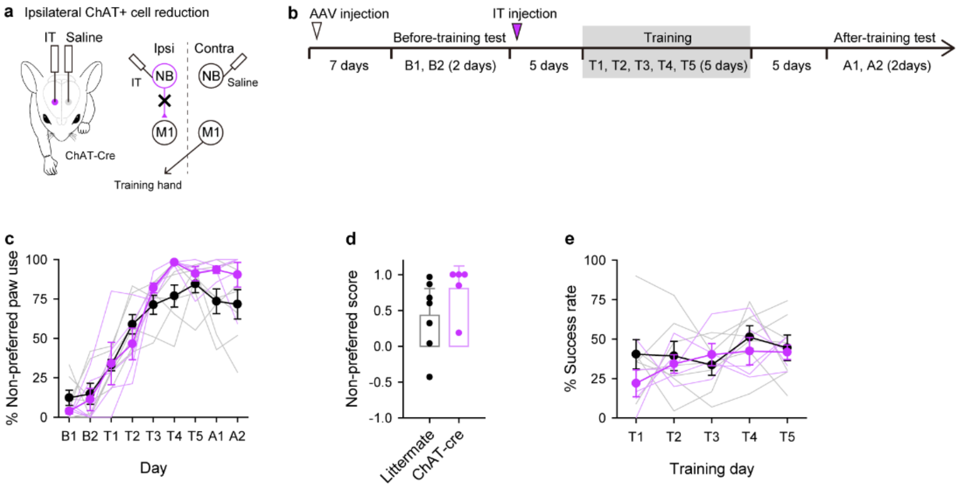


**Extended Data Fig. 2 | Ipsilateral NB ablation does not affect paw switching.**

**a-b**, Schematic and experimental timeline. **c**, Transition in non-preferred hand use in ChAT-Cre mice and littermate controls. Littermate controlwere from the same cohort used in Figure 2**c–i** (*n* = 7 littermates, 4 ChAT-cre mice, *P* = 0.067, two-way ANOVA. **d**, Non-preferred score on day 9 (session A2; *P* = 0.18, Student's *t*-test). **e**, Success rates during training (*P* = 0.27, two-way ANOVA).

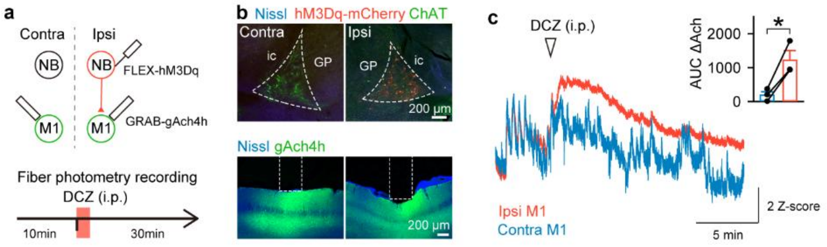


**Extended Data Fig. 3 | Unilateral DREADD-mediated activation of cholinergic neurons, monitored by fiber photometry.**

**a**, Schematic and experimental timeline for fiber photometry recording. **b**, Representative images showing the NB contralateral and ipsilateral to the hM3Dq expression site (top) and fiber insertion site in M1 (bottom). **c**, Representative traces of gAch4h signals from M1 ipsilateral and contralateral to the activation side. The white arrowhead indicates the timing of intraperitoneal injection of DCZ. Strength of gAch4h signals in response to DCZ in the contralateral and ipsilateral M1 ($n$ = 3 mice, *$P$ = 0.037, paired $t$-test).
